# CANDI: self-supervised, confidence-aware denoising imputation of genomic data

**DOI:** 10.1101/2025.01.23.634626

**Authors:** Mehdi Foroozandeh, Abdul Rahman Diab, Maxwell Libbrecht

## Abstract

Large-scale epigenomic datasets such as histone modifications and DNA accessibility have greatly advanced our understanding of genomic function. However, these measurements often suffer from noise, batch effects and irreproducibility. Epigenome imputation has emerged as a promising solution to these challenges. These methods integrate patterns across experiments, cell types, and genomic loci to predict the results of experiments, yielding predictions that often surpass observed data in quality. Thus, researchers increasingly leverage imputation for denoising data prior to downstream analysis.

However, existing methods for imputation-based denoising have significant limitations. Here, we propose CANDI (Confidence-Aware Neural Denoising Imputer), a method for epigenome imputation that (1) predicts raw counts and handles experiment-specific covariates such as sequencing depth, (2) can (optionally) incorporate information from a low-quality existing experiment when predicting a target without retraining, and (3) outputs a calibrated measure of uncertainty. This approach is enabled using a Transformer model with self-supervised learning (SSL) training.

## 1 Introduction

In recent years, the availability of large-scale functional genomic data such as histone modifications and DNA accessibility has provided unprecedented opportunities to understand the functional roles of diverse genomic loci. However, a major confounding factor is that measurements obtained using sequencing methods often suffer from various sources of noise, including batch effects, technical variability, and biological heterogeneity [1]–[6].

A promising approach for addressing issues of noise is epigenome imputation. Epigenome imputation methods aim to predict the output of a functional genomics experiment. Due to the high cost and complexity of profiling every possible assay in all relevant cell and tissue types, researchers have turned to computational methods to predict missing data, including ChromImpute, Avocado, eDICE and others [3], [7]–[10].

Epigenome imputation methods were originally designed to predict unperformed assays, but researchers have shown that imputed data often has better properties than observed data, even when such observed data sets are available. By integrating patterns across experiments, cell types, and genomic loci, imputation models average out noise distilling consistent and biologically meaningful signals into less noisy predictions [7]. Thus, researchers frequently apply imputation for denoising by re-imputing each assay before inputting the assay into downstream analysis.

However, existing approaches for imputation-based denoising have significant limitations. Most significantly, all existing imputation methods operate on idealized processed signal (for example, fold enrichment over control). They assume that this processing removes all batch effects and results in an idealized “signal strength.” This issue jeopardized the recent ENCODE Imputation Challenge, which aimed to evaluate epigenome imputation methods comprehensively. Upon receiving entries from all participants, the organizers found that, due to subtle differences between the train and test sets, a simple baseline outperformed most of the entrants.

Furthermore, to denoise a given experiment using current approaches, one must typically re-train the model without that particular experiment and then impute it *de novo*. This usually would require re-training the underlying machine learning model; thus, the only existing method that can be applied for denoising in practice is ChromImpute, whose learning architecture allows this process to be performed without re-training. This existing strategy also entirely disregards the target assay, which could provide valuable information towards denoising.

To address these issues, we propose CANDI (Confidence-Aware Neural Denoising Imputer), a method for epigenome imputation that (1) predicts raw counts and handles experiment-specific covariates such as sequencing depth, (2) can (optionally) incorporate information from a lowquality existing experiment when predicting a target without retraining, and (3) outputs a calibrated measure of uncertainty. This approach is enabled using self-supervised learning (SSL) with a Transformer model, a paradigm that capitalizes on large amounts of unlabeled data by corrupting and then reconstructing subsets of the input to learn without explicit labels. This strategy enables zero-shot imputation and denoising, allowing models to generalize to new cell types without retraining.

## 2 Results

### 2.1 Self-supervised, confidence-aware denoising imputation of genomic data

We propose CANDI, a method for epigenome imputation. Briefly, CANDI works as follows (Methods): For a given 30kb locus, CANDI takes as input (1) observed epigenomic data sets for the locus in the target sample along, (2) four experimental covariates for each observed assay, and (3) the DNA sequence of the locus. It also takes as input the experimental covariates of the desired outputs. CANDI outputs predicted epigenomic data sets for the given locus and sample in two formats: (1) raw read count and (2) continuous signal in log Poisson p-value units. Both outputs are given in the form of a probability distribution: a negative binomial distribution over raw reads and a Gaussian distribution over continuous signal.

CANDI consists of a neural network model that includes convolutional, Transformer and deconvolutional layers (Methods). We trained CANDI using a self-supervised learning approach by optimizing two objectives. First, we masked some fraction of training tracks and asked CANDI to predict these masked tracks. Second, we simulated low-quality data by downsampling reads from training tracks and asked CANDI to predict the original data from low-quality data.

### 2.2 CANDI accurately imputes missing epigenetic signals

To evaluate the performance of CANDI, we first assessed its ability to accurately impute missing epigenomic signals across diverse cell types and experimental assays. A genome-wide evaluation demonstrated strong performance across a wide range of epigenetic marks and cell types. In particular, CANDI achieved a high correlation between predicted and experimental signals for active regulatory marks such as H3K27ac and H3K4me3, with Pearson correlations exceeding 0.8 in many cell types, consistent with previous imputation methods (Figure 2C). CANDI effectively distinguishes between foreground and background signals. For instance, we analyzed H3K4me3, an active regulatory mark associated with promoter regions. The imputation performance was evaluated within ±200 bp of transcription start sites (TSS) on chromosome 21 of neural progenitor cells (Figure 2A, B). CANDI takes raw read counts and experimental covariates as input and produces both imputed raw counts and processed signals as output. We observe that CANDI performs better at predicting accurate read counts compared to processed signals (Fig 2C).

**Figure 1.**
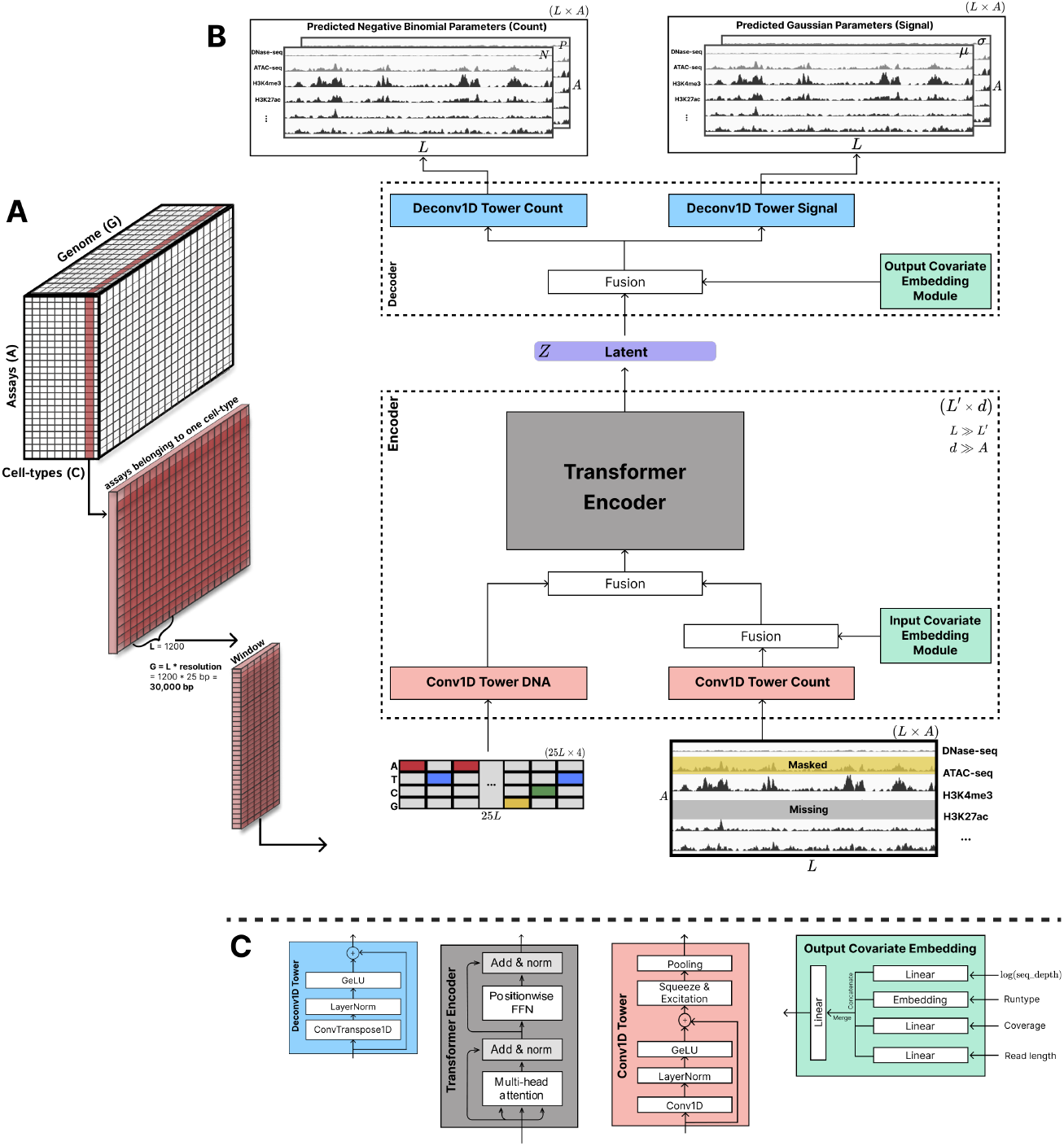
CANDI architecture overview. (A) Input data organization showing the three-dimensional structure of epigenomic data across assays, genomic positions, and cell types. (B) Model architecture consisting of encoder and decoder components. The encoder processes DNA sequence and epigenomic count data through parallel Conv1D towers, integrates experimental covariates, and generates a latent representation using a transformer encoder. The decoder predicts both count data (negative binomial parameters) and signal values (Gaussian parameters) for each assay and position. (C) Detailed architecture of key model components including Conv1D towers, transformer encoder blocks, and covariate embedding modules.

**Figure 2.**
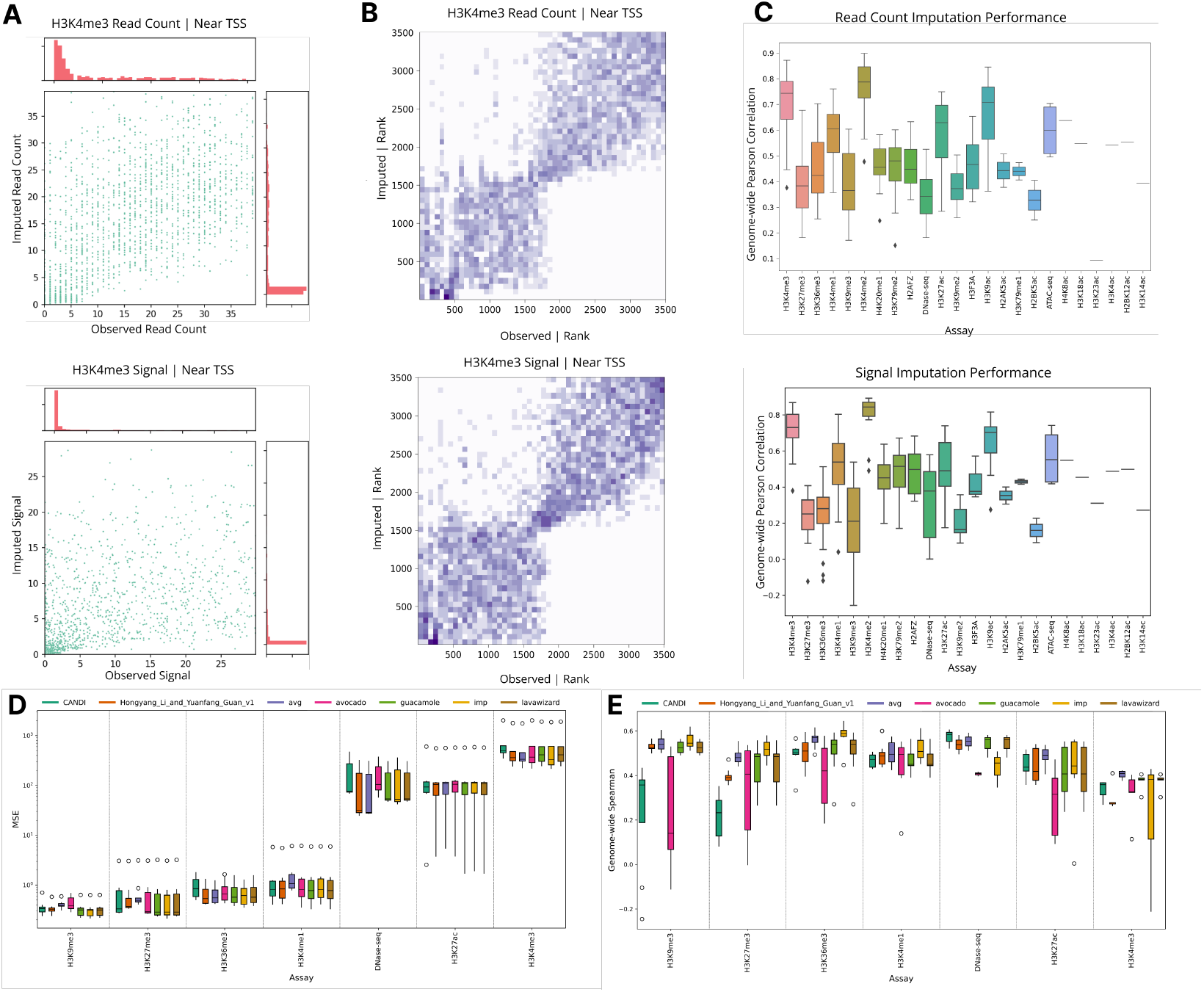
Evaluation of CANDI’s imputation performance. (A) Scatter plots comparing observed versus imputed H3K4me3 values within ±200bp of Transcription start site (TSS) regions on chromosome 21 in neural progenitor cells. The top panel shows read counts, while the bottom panel shows signal values. Marginal distributions are shown along each axis. Imputed read count is mean of the predicted negative binomial distribution. (B) Rank-based density heatmaps of the same H3K4me3 data, showing the relationship between observed and imputed ranks for both read counts (top) and signal values (bottom). The color intensity, presented on a log scale, indicates the density of points, with darker colors representing higher densities. (C) Boxplots displaying the genome-wide Pearson correlation coefficients for read count (top) and signal (bottom) predictions across different assays in the extended dataset test set, demonstrating CANDI’s performance across various epigenomic marks. (D-E) Comparative performance evaluation of CANDI against baseline methods and winners of the ENCODE Imputation Challenge, showing (D) Mean Squared Error (MSE) and (E) Spearman rank correlation metrics across different assays. The comparison includes Average (naive baseline), Avocado, and other top-performing methods from the challenge.

We benchmarked CANDI on the ENCODE Imputation Challenge (EIC) dataset [3], a standardized benchmark for evaluating epigenomic imputation methods. We compared CANDI-predicted signals with the following baseline and challenge-winning methods: (1) average (naive baseline), (2) Avocado [8], (3) Guacamole, (4) Lavawizard, (5) imp (an earlier version of eDICE [10] that participated in EIC), and (6) Hongyang Li and Yuanfang Guan v1. For this comparison, we followed the train-test split of the EIC dataset. CANDI achieved performance comparable to these methods in terms of Spearman rank correlation and mean squared error (Figure 2D, E).

Unlike other methods, CANDI performs imputation solely based on the combinatorial patterns of the available assays in a given cell type. It does not leverage information from other cell types and relies entirely on patterns learned by the model. At test time, CANDI has access only to the DNA sequence and observed assays of the target cell type. It does not receive explicit information about genomic positions or cell types, requiring the model to infer generalizable patterns rather than memorizing position- or cell type-specific signals. This design choice presents a more challenging problem compared to methods that explicitly leverage cell type or positional metadata. However, it encourages learning robust biological relationships and enables efficient zero-shot imputation on new cell types using patterns learned during self-supervised pretraining.

To avoid memorization bias, especially regarding DNA sequences, we excluded chromosome 21 from the CANDI training dataset and tested exclusively on this chromosome. While our benchmark comparisons followed the exact train-test split of the EIC dataset (Figure 2D, E), we adopted a different strategy for the extended dataset to simulate the more challenging cell type-agnostic scenario (Figure 2A,B,C). In the EIC train-test split, a subset of assays within a given cell type is reserved for testing or validation. For example, from 10 available assays in a given cell type, 7 may be used for training, and 3 reserved for testing. This approach assumes cell type-specific embeddings, where models must know the cell type at test time to impute the reserved assays. In contrast, our approach reserves entire cell types for training, testing, and validation. This ensures that at test time, the model encounters completely new cell types with only a few available assays, better mimicking real-world imputation. By doing so, we enforce a cell type-agnostic framework, requiring CANDI to generalize across unseen cell types based solely on learned patterns.

### 2.3 CANDI provides calibrated aleatoric uncertainty estimates

CANDI predicts probability distributions rather than single values, enabling it to capture the uncertainty in its predictions. Specifically, it captures aleatoric uncertainty, the uncertainty caused inherent randomness in experimental data generation caused by biological variation, technical noise, and experimental biases. This probabilistic approach provides insight into the variability of predictions due to factors intrinsic to the data. For each prediction, CANDI generates confidence intervals around the median value which allows for soft prediction of observed signals i.e. even if the expected value of the predicted distribution does not exactly match the observed value, it falls within the predicted confidence interval (Fig 3A, B)

**Figure 3.**
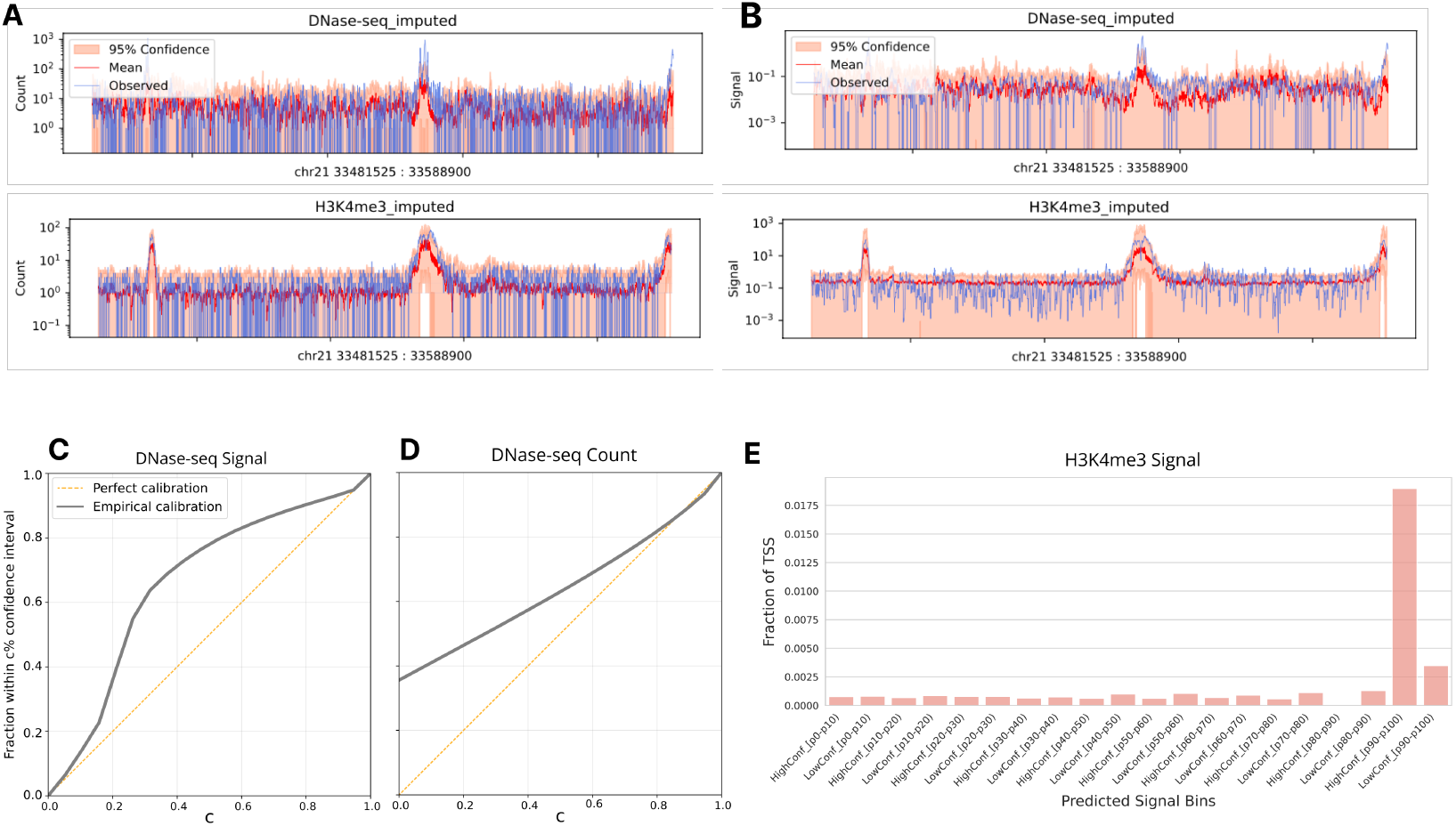
Uncertainty-aware imputation of epigenomic signals demonstrates accurate predictions with calibrated confidence estimates. A-B) Imputed DNase-seq (top) and H3K4me3 (bottom) signals around a housekeeping gene locus (GART) on chromosome 21 in neural progenitor cells, showing both read counts (A) and signal values (B). Red lines represent predicted mean values from Gaussian (signal) and negative binomial (count) distributions, with pink shading indicating 95% confidence intervals. Blue lines show observed experimental values. Y-axes are in log scale. C-D) Calibration plots for DNase-seq predictions in neural progenitor cells (chromosome 21) for both signal values and read counts. The horizontal axis shows confidence level (c), and vertical axis shows the empirical fraction of observations falling within the c% confidence interval. Gray lines above the diagonal indicate conservative uncertainty estimates. E) High-confidence imputed H3K4me3 signals in the top 10% predicted signal bins show significantly greater overlap with TSS compared to low-confidence predictions or low-signal regions.

**Figure 4.**
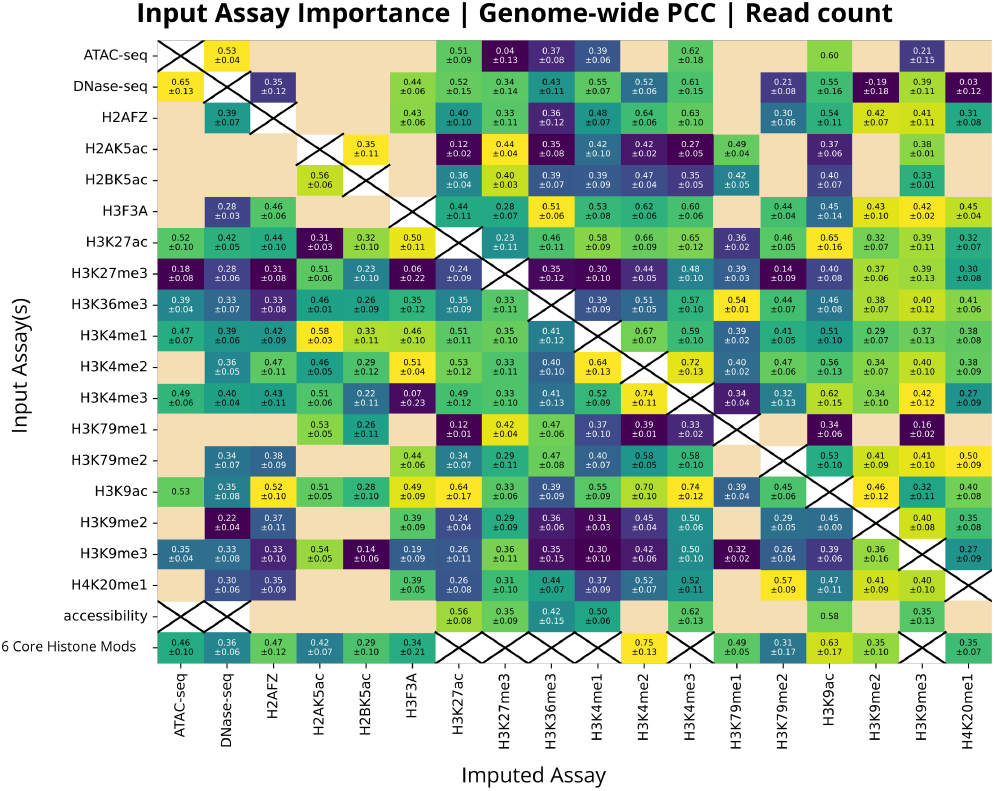
Input assay importance analysis for CANDI’s imputation performance: The heatmap shows genome-wide Pearson correlation coefficients (PCC) between predicted and actual read counts across different input configurations. The vertical axis represents three input scenarios: single assay inputs, accessibility assays (ATAC-seq + DNase-seq), and six core histone marks (H3K4me3, H3K4me1, H3K27ac, H3K27me3, H3K9me3, and H3K36me3). The horizontal axis indicates the target assays being imputed. Cells marked with an “X” represent cases where the target assay is included as an input. Beige cells indicate unavailable data. The color scale is normalized for each column, with the brightest cell representing the best predictor for the corresponding target assay.

To evaluate the quality of CANDI’s uncertainty estimates, we evaluated their calibration. For a well-calibrated model, we expect that the observed values should fall within the predicted *C*% confidence interval *C*% of the time. Our analysis shows that CANDI’s confidence intervals have near-perfect calibration for high confidence levels (*C*≥ 0.9), while being conservatively biased for lower confidence levels (Fig 3C, D). This conservative bias is partly caused by the discrete nature of count data: we observe that the predicted median of read counts (negative binomial) exactly matches the observed value 38% of the time (Fig 3D), meaning that even a 0% confidence interval overlaps the observed value 38% of the time.

Quantifying uncertainty is important when using imputed tracks. For example, for imputed H3K4me3 signals, regions with high predicted signal (*>*90th percentile) and higher confidence show significantly greater overlap with TSS regions compared to those with low-confidence high signals or low-signal predictions (Fig 3E). That is, imputed signals with higher certainty are more likely to correspond to real biology.

### 2.4 CANDI predicts across various input experiment combinations

Because CANDI can evaluate the uncertainty in its predictions, researchers can use it to prioritize which assays are most needed for a given target cell type. Moreover, CANDI can do so without expensive re-training.

To evaluate which input assays are most informative for predicting a given target assay, we conducted a comprehensive analysis to understand CANDI’s predictive capabilities under different input scenarios. Our investigation focused on three key conditions: single-assay input, accessibility data (DNase and ATAC-seq), and six core histone modifications. This systematic approach helped identify the most informative input assays for predicting any given target assay.

Our analysis revealed that the predictive power of CANDI varies significantly depending on the available input assays. Most notably, we found strong predictive relationships between functionally related assays. For instance, DNA accessibility markers (DNase and ATAC-seq) showed high mutual predictability, demonstrating CANDI’s ability to leverage biological relationships in its predictions. Conversely, repressive marks such as H3K27me3 and H3K9me3 provided limited predictive value for active regulatory marks, reflecting the distinct biological roles of these modifications.

## 3 Discussion

Effective denoising imputation with CANDI promises to greatly improve all downstream applications of epigenomic data sets, including chromatin state annotation and predictive modeling. CANDI enables the creation of “supertracks” that simulate a complete data set with excellent experiment quality and uniform experimental parameters.

## 4 Methods

### 4.1 Data Collection and Processing

We trained and evaluated CANDI using two datasets.

#### ENCODE Imputation Challenge Dataset

We used data from the ENCODE Imputation Challenge (EIC), featuring 35 distinct assays measured across 50 biosamples. For each experiment, we obtained aligned sequencing reads (BAM files), signal p-values (BigWig files), and experimental covariates, including sequencing depth, coverage, read length, and run type (single- or paired-end). In the original ENCODE Imputation Challenge, the dataset contained only BigWig files of signal p-values. However, since we use read counts as input to our model, we obtained raw read files (BAM) corresponding to the same experiments and biosamples in the EIC. We followed the original train-validation-test split proposed by the EIC to ensure comparability with previous benchmarks.

#### Extended ENCODE Dataset

To have a more extensive dataset, we systematically collected data for all biosamples in the ENCODE database that contained at least one experiment from the 35 assays of interest. This initial collection yielded 3064 biosamples, forming a sparse 3064 *×*35 experiment availability matrix. To address this sparsity while maintaining biological relevance, we implemented a cell type-based merging strategy. This merging strategy significantly improved data density, increasing the median number of available assays per sample from 1 (per biosample) to 6 (per merged cell type). The final processed dataset consists of 361 merged cell types.

To simulate varying sequencing depths, we implemented read downsampling in BAM files. The Downsampling Factor (DSF) determines the fraction of reads retained: DSF 1 retains all reads, DSF 2 randomly samples 50% of reads, and DSF 4 randomly samples 25% of reads.

### 4.2 CANDI architecture

Let *A* = 35 represent the number of assay types, and let *G* = 30,000 bp and *R* = 25 bp denote the length of the genomic window and the resolution, respectively, such that the window contains *L* = *G/R* = 1200 genomic positions at 25 bp resolution. Each sample (i.e., one data instance in a batch) corresponds to epigenomic data from a single cell type, forming a matrix **M** of shape *A × L* for that cell type. We process multiple samples in a batch of size *N* = 50, where each sample has its own **M, S**, and covariates.

The model receives: **M** ∈ ℝ^*N ×A×L*^, **S** ∈ {0, 1*}*^*N ×*4*×G*^, **C**_*in*_ ∈ ℝ^*N ×A×*4^, **C**_*out*_ ∈ ℝ^*N ×A×*4^.

**Inputs**.

- Epigenomic Reads **M**: Each *M*_*a,l*_ is the read count for assay *a* at position *l*.
- DNA Sequence **S**: One-hot encoded nucleotides across the *G* = 30,000 bp region.
- Covariates **C**_*in*_ and **C**_*out*_: Four values per assay (log_2_(depth), type, read length, coverage).

**Outputs**. For each assay *a* at each position *l*, the model predicts parameters of probability distributions:

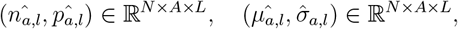

where 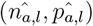 parameterize Negative Binomial distributions for raw counts for assay *a* at position *l* and 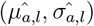 parameterize Gaussian distributions for processed signal p-values for the same assay at the same position.

**Model Overview**. The model consists of an encoder ℰ and a decoder 𝒟:

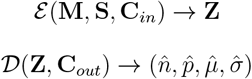

**Encoder** *ℰ*. The encoder integrates epigenomic data **M**, DNA sequence context **S**, and experimental covariates **C**_*in*_ into a latent representation **Z**.

#### 1. Epigenetic Signal Convolution Tower

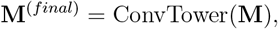

resulting in a transformed epigenomic representation 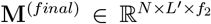,where 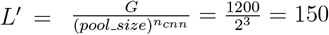 and

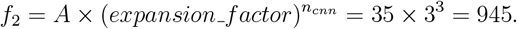

Here, *A* = 35 is the number of assays. This tower processes the epigenomic signals with *n*_*cnn*_ = 3 depth-wise convolution layers, each with *pool size* = 2, progressively reducing the spatial dimension from *L* = 1200 to *L*^*′*^ = 150 and expanding the feature dimension. The expansion factor 3 scales feature dimensions through the convolution towers. The convolution kernel size is 3 across all layers.

#### 2. DNA Sequence Convolution Tower

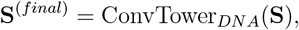

producing 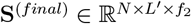,where

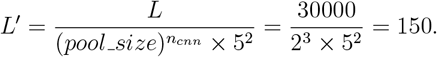

In addition to the *n*_*cnn*_ convolutional layers with *pool size* = 2, the DNA sequence convolution tower includes two additional layers at the end, each with *pool size* = 5. These final layers compensate for the difference in resolution between the epigenomic signals (25 bp resolution) and the DNA sequence (1 bp resolution), ensuring both modalities align at the same final context length *L*^*′*^. The convolution kernel size is 3 across all layers.

#### 3. Input Covariate Embedding

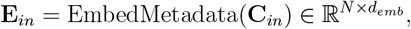

Here, *d*_*emb*_ = *A*, and **C**_*in*_ consists of four covariates per assay: one categorical (sequencing type i.e. single or pair end) and three continuous (log_2_(sequencing depth), read length, and coverage). Each continuous covariate is linearly transformed into an embedding, and the categorical covariate is mapped via a learned embedding. These four embeddings per assay are concatenated, normalized, and projected into a final *A*-dimensional vector per sample.

#### 4. Fusion with Epigenetic Signals

Concatenate **E**_*in*_ with **M**^(*final*)^ along the feature dimension and map back 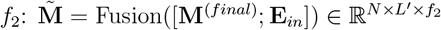.

#### 5. Fusion with DNA Features

Concatenate 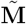 and **S**^(*final*)^ as 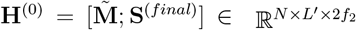, and transform it to 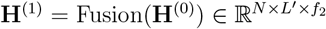.

#### 6. Transformer Encoder (Relative Positional Encoding)

Apply 4 transformer encoder layers with *n*_*head*_ = 9 attention heads and a relative positional encoding scheme **Z** = TransformerEncoder(**H**^(1)^), resulting in the latent representation 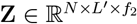.

**Decoder** *𝒟*. Given **Z** and **C**_*out*_, the decoder reconstructs the signal distributions:

1. **Output Covariate Embedding**:

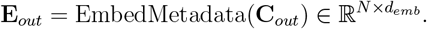
2. **Fusion with Latent Representation**:

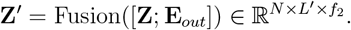
3. **Deconvolution Towers**: The latent representation 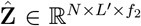 is passed to two separate decoder modules: one dedicated to reconstructing read counts, and another dedicated to predicting processed signal values. Each decoder employs a sequence of “DeconvTower” layers that invert the encoder’s pooling and feature expansion steps. Formally: **X**^(*rec*)^ = DeconvTower(**Z**^*′*^) ∈ ℝ^*N ×L×A*^, that results in a restored resolution along the genomic positions *L* and assays *A*.
4. **Distribution Layers**:

Each decoder’s reconstructed output is then passed through a corresponding distribution layer: 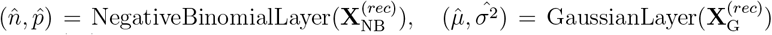,where 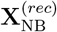 and 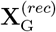 are the feature maps from the count and signal decoders, respectively.

In the **NegativeBinomialLayer**, the input **X**^(*rec*)^ ℝ^*N ×L×A*^ is passed through two separate linear transforms and layer normalizations. For each assay-position pair:

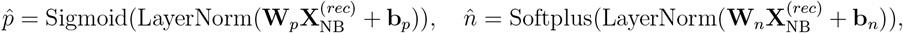

ensuring 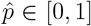 and 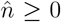.Thus, 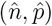 parameterize a negative binomial distribution for raw counts.

In the **GaussianLayer**, the input 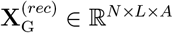 is likewise passed through linear and normalization layers:

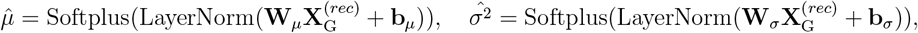

ensuring 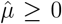 and 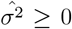.Thus, 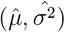 define a Gaussian distribution over processed signal values.

By producing 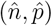 and 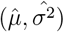 in this manner, CANDI outputs full probability distributions— rather than single-point estimates—allowing it to capture uncertainty inherent in epigenomic signals.

### 4.3 Learning objectives and training strategy

Our approach leverages self-supervised learning, where the model learns to reconstruct deliberately corrupted versions of its own input. We employ two complementary learning objectives:

1. **Masking:** Unlike conventional masking methods that mask positions, our approach masks entire features (assays). We randomly mask a subset of assays for each cell type and challenge the model to predict the masked assays from the remaining available ones at the same genomic loci. This task simulates the imputation scenario, where some experiments are missing and must be inferred from those that are present.
2. **Upsampling:** We artificially downsample selected assays, thereby reducing their effective sequencing depth, and train the model to predict the original higher-depth signals. By doing so, the model learns to enhance low-depth experiments, effectively “upsampling” them to higher-quality signals.

Formally, for each assay *a* and genomic position *l*, we have observed raw counts *y*_*l,a*,raw_ and processed signal values *y*_*l,a*,signal_.

We train CANDI by minimizing the following objective, which combines four negative log-likelihood loss functions:

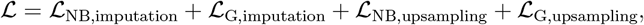

where:

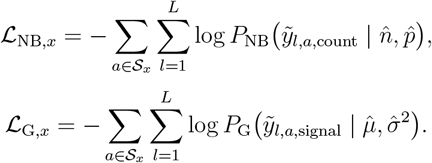

Here, *x*∈ {imputation, upsampling specifies} the task, and 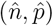 and 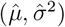 are the predicted parameters of the negative binomial and Gaussian distributions, respectively. The terms with tildes 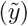 represent the original data before downsampling.

#### Training loci

To optimize model training, we focus on genomic regions containing candidate cis-regulatory elements (cCREs), which are enriched with informative and functionally relevant epigenomic signals.

We trained CANDI on 3,000 randomly selected, non-overlapping regions, each spanning 30kb and containing at least one cCRE. We excluded chromosome 21, reserving it exclusively for testing purposes. In total, CANDI was trained on 90 million base pairs of genomic sequence, representing approximately 3% of the human genome.

### 4.4 Imputation performance

To evaluate imputation performance, we compute the following genome-wide metrics: Mean Squared Error (MSE) for average squared differences, Pearson Correlation for linear relationships, and Spearman Correlation for monotonic relationships between observed and imputed values. These metrics demonstrate the model’s ability to impute genomic signals. The main CANDI model is trained on the full dataset, but to ensure comparability with the ENCODE Imputation Challenge (EIC) competitors, we trained a version of CANDI using the EIC dataset and their specified train-test split. All performance evaluations, including on EIC baseline and winning methods, were made on chromosome 21.

### 4.5 Uncertainty modeling

#### Confidence calibration

To evaluate the reliability of the model’s uncertainty estimates, we assess its calibration. Specifically, we measure the fraction of observed values that fall within the model’s predicted *C*% confidence interval for *C*∈ [0, 1]. A well-calibrated model should exhibit a linear relationship, with *C*% of observations lying within the *C*% confidence interval. This provides insight into how well the model quantifies its uncertainty.

#### Functional validation

As an additional biologically relevant assessment, we evaluate whether the model confidently predicts high H3K4me3 signal (a mark associated with promoters) near transcription start sites (TSS).

## Data and Code Availability

All codes, implementation details, and data collection procedures are available at” https://github.com/mehdiforoozandeh/EpiDenoise.

